# A detailed analysis of 16S rRNA gene sequencing and conventional PCR-based testing for the diagnosis of bacterial pathogens and discovery of novel bacteria

**DOI:** 10.1101/2024.06.26.600821

**Authors:** Mei-Na Li, Ting Wang, Nan Wang, Qiang Han, Xue-Ming You, Shuai Zhang, Cui-Cui Zhang, Yong-Qiang Shi, Pei-Zhuang Qiao, Cheng-Lian Man, Teng Feng, Yue-Yue Li, Zhuang Zhu, Ke-Ji Quan, Teng-Lin Xu, George Fei Zhang

**Affiliations:** International Joint Research Center for Microbiology and Infectious Diseases, Wohua Biotech, Binzhou, 256600, Shandong, China; Department of Basic and Forensic Medicine, North Sichuan Medical College, Nanchong, 637100, Sichuan, China

**Author notes:** Correspondence: George Fei Zhang, (Lead contact), Teng-Lin Xu, Ke-Ji Quan. These authors contributed equally to this work (Mei-Na Li, Ting Wang, Nan Wang).

**Keywords:** 16S rRNA gene sequencing, conventional polymerase chain reaction, diagnosis, bacterial community, *Avibacterium paragallinarum*, swollen head syndrome

## Abstract

This study represents the first analysis of the bacterial community in chickens affected by swollen head syndrome, utilizing 16S rRNA gene sequencing. Samples were obtained from clinical laying chickens and were examined for the presence of *Avibacterium paragallinarum* (APG) and *Ornithobacterium rhinotracheale* (ORT) using conventional polymerase chain reaction (PCR). From the samples, five APG-positive (APG) and APG-negative (N-APG) samples were chosen, along with five specific pathogen-free chickens, for 16S rRNA gene sequencing. Results showed that APG and ORT were widely detected in the chicken samples with swollen head syndrome (SHS, 9/10), while APG was detected in all five specific pathogen-free (SPF) samples. In contrast, conventional PCR sensitivity was found to be inadequate for diagnosis, with only 35.7% (5/14) and 11.1% (1/9) sensitivity for APG and ORT, respectively, based on 16S rRNA gene sequencing data. Furthermore, 16S rRNA gene sequencing was able to quantify the bacteria in the samples, revealing that the relative abundance of APG in the APG group ranged from 2.7% to 81.3%, while the relative abundance of APG in the N-APG group ranged from 0.1% to 21.0%. Notably, a low level of APG was also detected in all 5 SPF samples. The study also identified a significant number of animal and human common bacterial pathogens, including but not limited to *Gallibacterium anatis*, *Riemerella columbina*, *Enterococcus cecorum, Mycoplasma synoviae*, *Helicobacter hepaticus*, and *Staphylococcus lentus*. In conclusion, 16S rRNA gene sequencing is a valuable tool for bacterial pathogen diagnosis and the discovery of novel bacterial pathogens, while conventional PCR is not reliable for diagnosis.

## 1 Introduction

Polymerase chain reaction (PCR) is a widely used method for diagnosing infectious diseases in humans and animals. However, conventional PCR may not be able to detect samples with a low number of copies (He et al., 1994). In addition to real-time and digital PCR, 16S rRNA sequencing is also a useful tool for bacterial identification and disease diagnosis (Fida et al., 2021; Rampini et al., 2011).

Swollen head syndrome (SHS), a condition that affects broilers, broiler breeders, and layers, is a widespread problem in many countries around the world. First reported in 1984 in South Africa, SHS has since been identified in numerous other countries, including Spain, France, the United Kingdom, the Netherlands, Canada, Israel, Japan, Iraq, and China (Al-Hasan et al., 2021; Hafez and Löhren, 1990; Nakamura et al., 1997). Several causative agents, including *Avibacterium paragallinarum* (APG), *Rhinotracheitis* virus, *Escherichia coli*, *Ornithobacterium rhinotracheale* (ORT), and *Metapneumovirus*, have been identified thus far (Abdelmoez et al., 2019; Al-Hasan et al., 2022; Al-Hasan *et al*., 2021; Droual and Woolcock, 1994). Identifying these agents is crucial in preventing the spread of SHS.

APG is a significant causative agent to chicken disease, as it causes infectious coryza (IC), an acute upper respiratory disease (Gallardo et al., 2020; Paudel et al., 2017). IC is characterized by mucous nasal discharge, facial swelling, and conjunctivitis (Balouria et al., 2019; Xu et al., 2019). Moreover, IC is responsible for decreased egg production (10%-40%) and increased unthrifty chickens (Heuvelink et al., 2018). Additionally, IC can have negative consequences for broiler chickens, as the reported IC cases in California resulted in increased mortality (8%-15%), leading to significant financial losses for the chicken industry (Crispo et al., 2019). Another bacterial pathogen identified in SHS is ORT, which is reported globally (Al-Hasan *et al*., 2021; Barbosa et al., 2019). ORT infection causes respiratory symptoms, growth retardation, reduced egg production, and mortality, resulting in economic losses for the poultry industry. Specifically, the respiratory symptoms of ORT infection consist of tracheitis, pericarditis, sinusitis, and exudative pneumonia, with fibrin purulent lesions and often unilateral pneumonia (Barbosa *et al*., 2019).

Co-infection with other bacteria or viruses can often affect the pathogenicity and persistence of APG in the host, and the environmental conditions also play a crucial role. Various studies have highlighted the importance of early diagnosis (Ali et al., 2022a; Ali et al., 2024; Ali et al., 2022b; Zhang et al., 2021) and vaccination (Quan et al., 2018; Zhang et al., 2015; Zhang et al., 2018a; Zhang et al., 2018b; Zhang et al., 2022; Zhang et al., 2023; Zhu et al., 2017) as effective tools in combating IC and preventing the infection of other pathogens.

The human and animal bodies are inhabited by a diverse community of microorganisms, such as archaea, bacteria, fungi, and viruses (Libertucci and Young, 2019). The state of health and disease depends on the interplay between the host’s immune responses, the native microbiota, and possible pathogens. Maintaining a balanced bacterial community can help prevent infections by providing colonization resistance (Libertucci and Young, 2019). Analyzing bacterial communities has the potential to pinpoint the pathogen responsible for an infection.

Recent studies have documented the bacterial community in healthy and infected human hosts. However, the microbiota of chickens affected by infectious coryza remains unexplored. To address this gap, we conducted a study in which we analyzed samples from chickens with swollen head syndrome for APG and ORT using PCR. Subsequently, we examined the bacterial community of the chicken infraorbital sinus through 16S rRNA sequencing and compared it to that of SPF chickens.

## 2 Materials and Methods

### 2.1 Study design and sample collection

This study compared the effectiveness of conventional PCR and 16S rRNA sequencing in detecting pathogens. Aseptic clinical samples (swabs) were collected from chicken infraorbital sinuses with SHS and categorized into APG positive (APG, 5 samples) and APG negative (N-APG, 5 samples) (Table 1). To identify and type APG, a multiplex PCR was carried out as described previously (Table 2) (Sakamoto et al., 2012). SPF chicken samples (Mock group) were also tested for APG, and all samples were screened for ORT using PCR. Additionally, bacteria isolation and 16S rRNA sequencing were performed on all collected samples. The chickens with SHS were sourced from various locations in China, while Wohua Biotech Animal Center provided the SPF chickens. All experiments were conducted with the approval of the Committee on the Ethics of Animal Experiments of Wohua Biotech company.

**Table 1.**
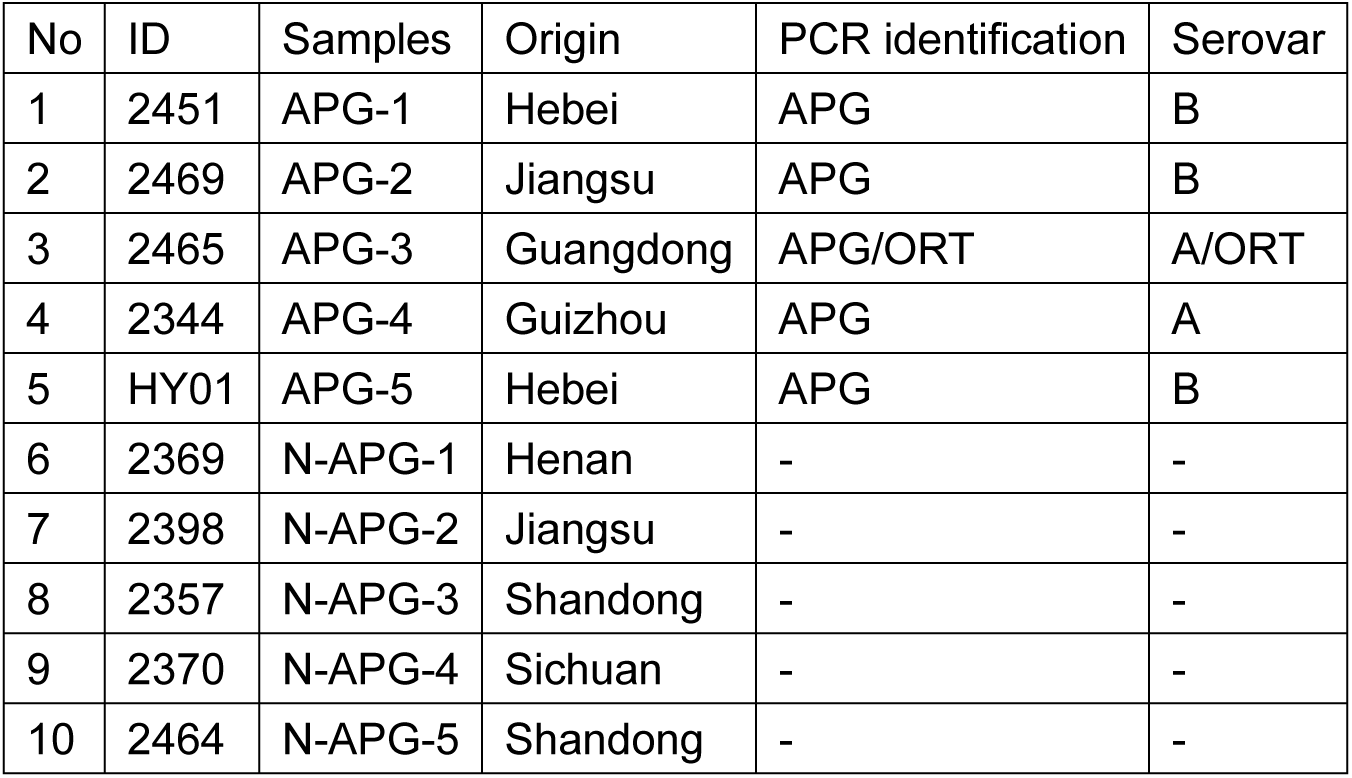
Samples in this study.

**Table 2.**
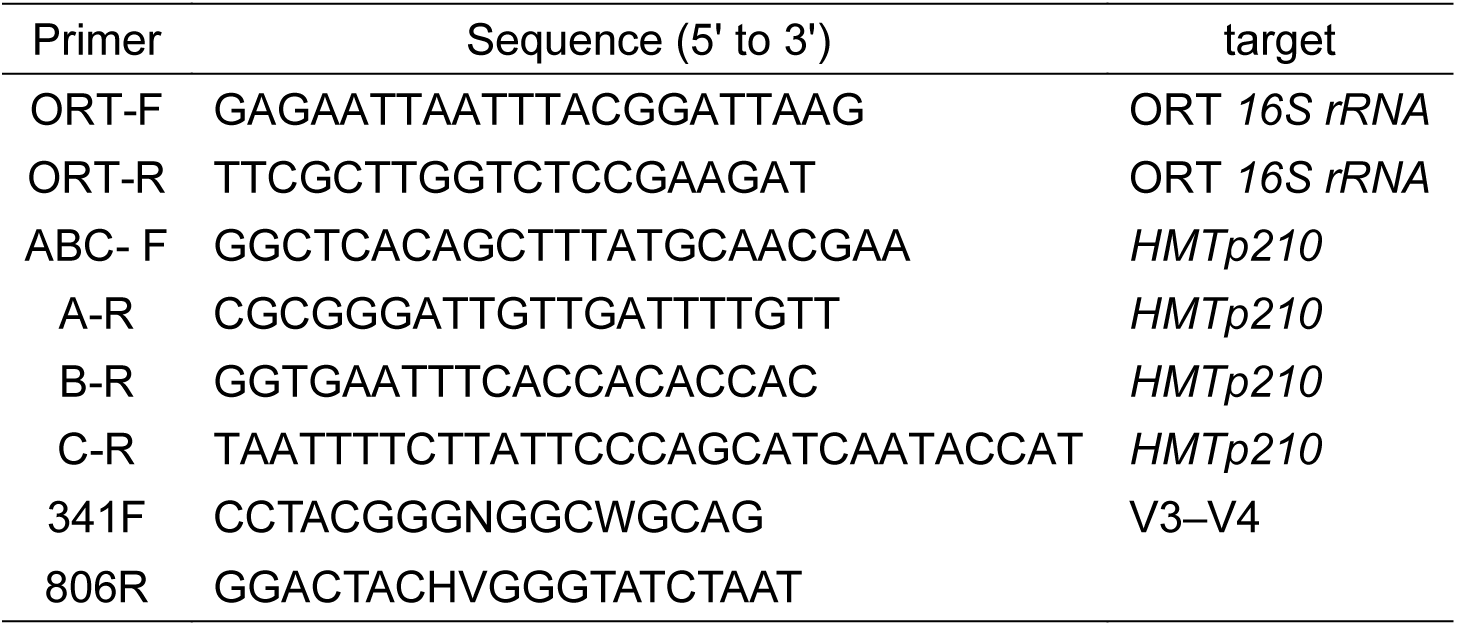
Primers used in this study.

### 2.2 Isolation and identification of *Avibacterium paragallinarum*

Chicken infraorbital sinus swabs were collected using the previously described method (Blackall and Soriano-Vargas, 2020). Briefly, the area under the eyes was cauterized with a hot iron spatula, followed by an incision in the sinus cavity using sterile scissors. A sterile cotton swab was then inserted deep into the sinus cavity for sampling. Swab samples were collected for PCR testing, 16S rRNA sequencing, and bacteria isolation. Samples were boiled in ultrapure water to serve as a PCR template for identifying APG and ORT. Swab samples were inoculated onto Tryptic Soy Agar (TSA) supplemented with 5% chicken serum and 50 μg/mL nicotinamide adenine dinucleotide (NAD). The bacteria were cultured overnight at 37 °C with 5% CO_2_, and their colonies were identified by PCR using specific primers of APG and ORT (Table 2). The samples were then tested with multiple PCR to amplify the *HMTp210* gene of serotypes A, B, and C and the *16S rRNA* gene of ORT (Table 2). DNA from serotypes A, B, and C of APG strains were used as templates for PCR.

### 2.3 DNA extraction, PCR amplification, and 16S rRNA sequencing

We collected ten samples (infraorbital sinus swabs) from chickens with SHS and five samples from SPF chickens (Mock) for 16S rRNA sequencing. We extracted the microbial DNA from each sample using the HiPure DNA Kits (Magen, China) following the product introduction. We quantified the DNA extractions using ultraviolet spectroscopy and amplified the V3–V4 domain of bacterial 16S rRNA genes by PCR (95 °C for 5 min, followed by 30 cycles at 95 °C for 1 min, 60 °C for 1 min, and 72 °C for 1 min, and a final extension at 72 °C for 7 min) using universal primers (Table 2). A 50 μL PCR mixture contained 10 μL of 5X Reaction Buffer, 10 μL of 5X High GC Enhancer, 1.5 μL of 2.5 mM dNTPs, 1.5 μL of each primer (10 μM), 0.2 μL of High-Fidelity DNA Polymerase, and 50 ng of template DNA (sequencing library preparation). We analyzed the amplicons using 2% agarose gels and purified them using the AxyPrep DNA Gel Extraction Kit (Axygen Biosciences, USA) following the manufacturer’s instructions. We normalized, pooled, and sequenced the purified amplicons on the Illumina NovaSeq 6000 Sequencing System according to standard protocols.

### 2.4 Bioinformatic analysis

We conducted 16S rRNA sequencing analysis as follows. Quality control, clustering, and high-throughput search were performed using Usearch, a robust sequence analysis software (http://www.drive5.com/usearch/). To ensure high-quality clean reads, raw reads were further filtered and assembled using FASTP (Chen et al., 2018). Clean tags were then clustered into operational taxonomic units (OTUs) of ≥ 97% similarity using UPARSE (Edgar, 2013). To classify representative OTU sequences into organisms, we used a naïve Bayesian model with an RDP classifier based on the SILVA rRNA database (https://www.arb-silva.de/) (Wang et al., 2007). Our analysis included species composition, indicator species, alpha diversity, beta diversity, and function analysis, which were all conducted using Omicsmart (http://www.omicsmart.com). The abundance statistics of domain, Phylum, Class, Order, family, genus, and species taxonomy were visualized in a stacked bar plot using Omicsmart. We also generated a network of correlation coefficients using Omicsmart. The raw sequencing reads were deposited into the Sequence Read Archive (SRA) database of NCBI with an accession number PRJNA1080319.

### 2.5 Statistical analysis

All statistical analyses were conducted using GraphPad Prism software. One-way ANOVA with Turkey’s multiple comparison test was employed to determine significant differences among groups. A P value of less than 0.05 was considered statistically significant. The results are presented as means with standard error of the mean (SEM) for each group.

## 3 Results

### 3.1 Bacterial identification by 16S rRNA sequencing and conventional PCR

PCR was used to examine Infraorbital sinus samples of SPF chicken and chicken with SHS for APG and ORT. Positive controls for APG serotypes A, B, and C were set in the PCR. A total of 5 APG positive samples and 5 APG negative samples of chicken with SHS were identified and collected for 16S rRNA sequencing analysis (Figure 1). Three serotype B (APG-1, APG-2, and APG-5) and 2 serotype A (APG-3 and APG-4) APGs were identified and isolated. One of the five APG-positive samples was also identified as ORT-positive (Figure 1A). In contrast, samples from SPF chicken were determined to be negative for both APG and ORT (Figure 1G). This study compared conventional PCR with 16S rRNA sequencing for identifying bacteria. Positive bands on the gel were considered as positive. Results showed that 14 out of 15 samples were identified as APG-positive by 16S rRNA sequencing, while only 5 out of 15 were identified by conventional PCR. Compared to 16S rRNA sequencing, the sensitivity of conventional PCR for APG detection is 35.7% (5/14). For ORT, only 1 out of 15 samples was identified as positive by conventional PCR, while nine out of fifteen were identified as positive by 16S rRNA sequencing (Figure 1, Supplemental Figure 1). Compared to 16S rRNA sequencing, the sensitivity of conventional PCR for ORT detection is 11.1% (1/9). The study concludes that 16S rRNA sequencing has a much higher sensitivity compared to conventional PCR.

**Figure 1.**
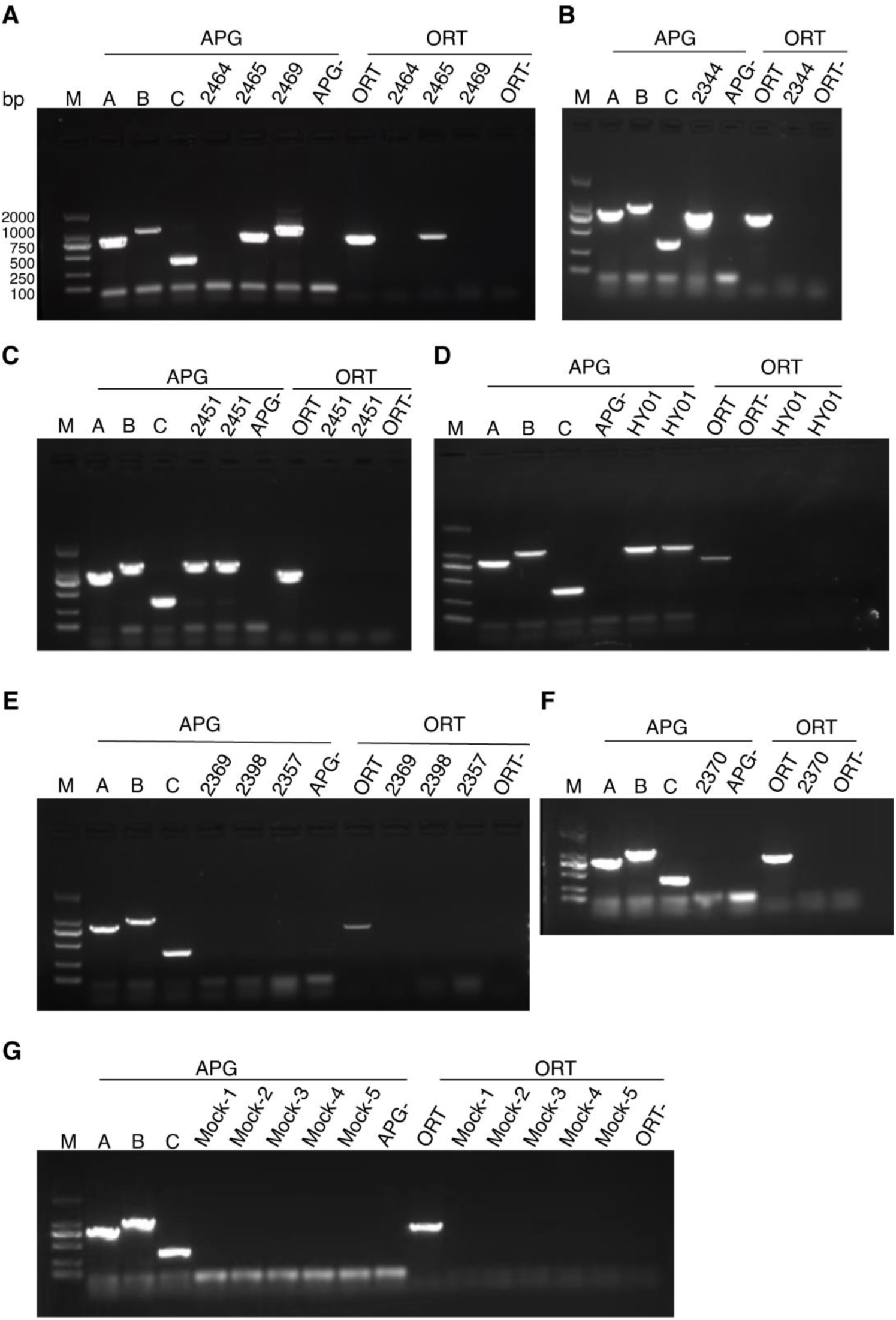
The PCR method was employed to identify APG and ORT in samples from commercial layer and SPF chickens (A-G). Lane M denotes the molecular weight standard, while samples A, B, and C serve as positive controls for APG serotypes A, B, and C. In total, ten clinical samples (2451, 2469, 2465, 2344, HY01, 2369, 2398, 2357, 2370, and 2464) and five SPF chicken samples underwent conventional PCR testing for APG and ORT.

### 3.2 Species composition analysis

Figure 2G lists the top 10 species, with APG being the most abundant in the APG-positive group. APG was found to be the predominant bacteria in most of the samples from the APG group, with abundance percentages of 2.69%, 81.26%, 65.50%, 76.58%, and 43.76% (APG-1 to APG-5), respectively. Although APG was not detected in the APG-negative group by PCR, sequencing data showed a relatively high abundance in those samples. Specifically, N-APG-1, N-APG-2, N-APG-3, and N-APG-4 showed abundance percentages of 20.96%, 1.18%, 1.79%, and 0.062%, respectively. It is worth noting that APG was found not only in infected chickens but also in SPF chickens (Table S1). The abundance of APG in the APG group was significantly higher than that of the Mock (P=0.0036, Figure 3) and N-APG (P=0.0113, Figure 3) groups. PCR detected ORT in one of the selected samples (Figure 1), with a higher abundance of ORT in that sample found by 16S rRNA sequencing (0.65%, Table S1). ORT was also widely detected in samples from chickens with SHS by 16S rRNA sequencing, although at too low an abundance to be detected by PCR (Figure 1, Table S1).

**Figure 2.**
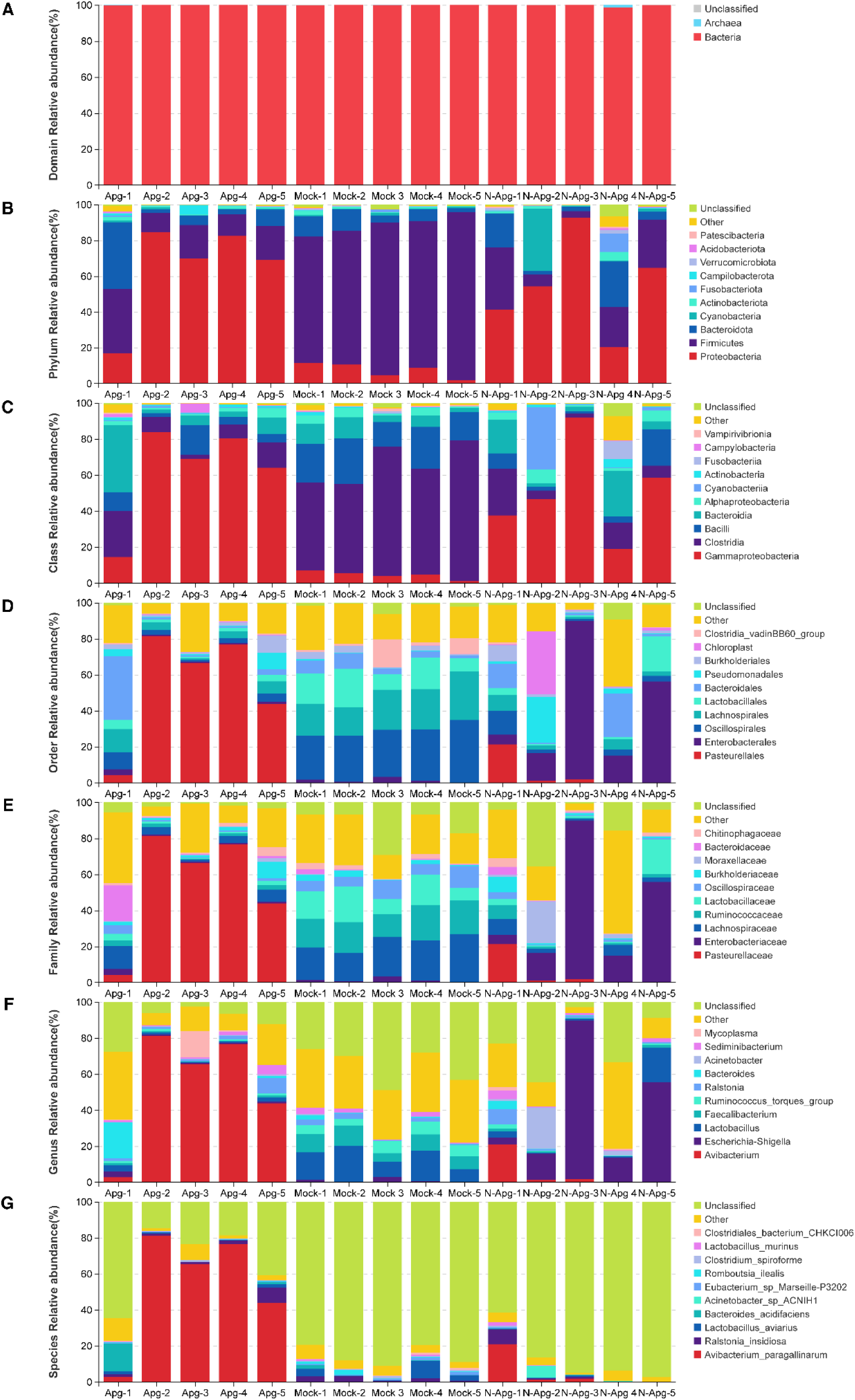
Analysis of the composition of bacterial communities. Through 16S rRNA amplicon sequencing, we identified the microbiota inhabiting the infraorbital sinus of 15 chickens. We analyzed the relative abundances of domain, phylum, class, order, family, genus, and species, and displayed the top ten (A-G).

**Figure 3.**
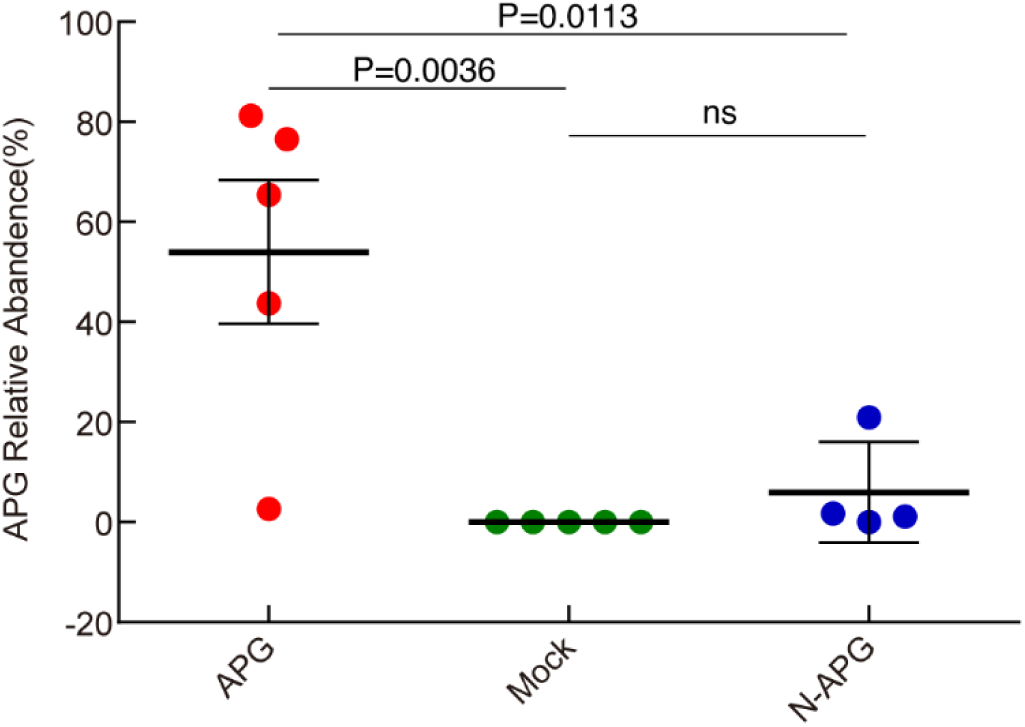
Analysis of the relative abundance of APG in APG, N-APG, and Mock groups. Compare the relative abundance of APG across different groups, and note that it is significantly higher than that of Mock and N-APG groups (A, B, and C).

### 3.3 Indicator species analysis

To compare the species in the Mock, APG, and N-APG groups, a Venn analysis was conducted using the R project Venn Diagram package (version 1.6.16) to identify unique and common species (Figure 4). The results showed that 1179, 1243, and 1885 OTUs were identified in Mock, APG, and N-APG, respectively (Figure 4). As illustrated in Figure 4A, 409 OTUs were shared by all groups. In terms of biological classification, from phylum to species analysis (Figure 4B-G), APG and N-APG exhibited a larger number of unique bacteria compared to the Mock group. To calculate the comparison of species between the different groups, Welch’s t-test in the R project Vegan package (version 2.5.3, Supplemental Figure 2) was used. The results showed that the abundance of APG in the APG group is significantly higher than that in the Mock group (P=0.02, Supplemental Figure 2). Furthermore, significant differences in the abundance of *Eubacterium sp Marseille-P3202*, *bacterium ic1379*, *Akkermansia muciniphila*, *Alistipes Sp CHKCI003*, *bacterium ic1296*, *Ralstonia pickettii*, *Coriobacteriaceae bacterium CHKCI002*, *Shewanella SP FDAARGOS 354* between Mock and APG group were also detected (Supplemental Figure 2). The Mock group exhibited a significantly higher abundance of *Eubacterium sp Marseille-P3202*, *Alistipes Sp CHKCI003*, bacterium ic1296, and *Coriobacteriaceae bacterium CHKCI002* than the N-APG group. Meanwhile, the APG group showed a significantly higher abundance of *Avibacterium paragallinarum*, *Alistipes inops*, and *Bacteroides sp Smarlab 3302398* than the N-APG group (Supplemental Figure 2).

**Figure 4.**
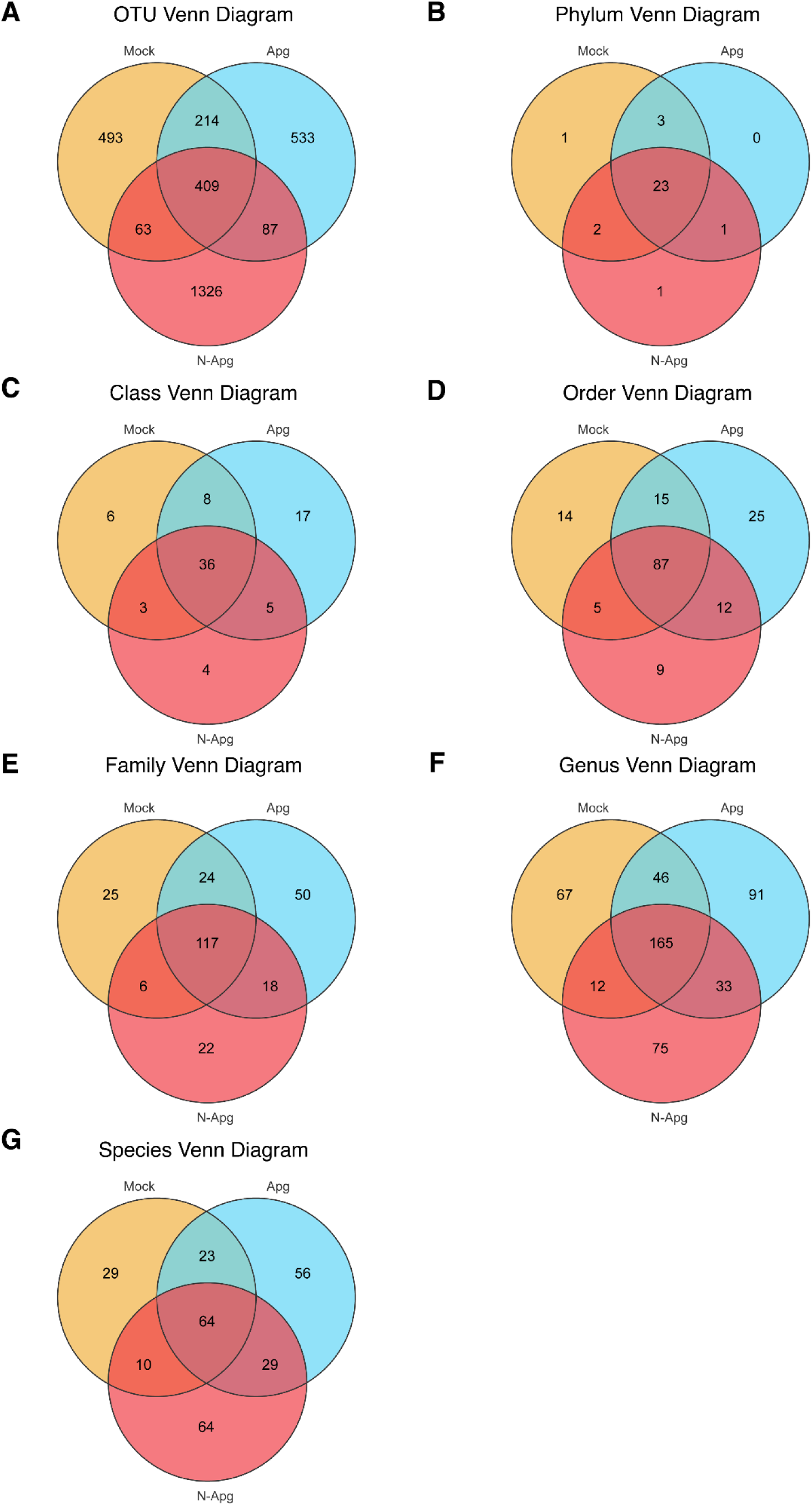
Indicator species analysis. Venn diagrams are used to illustrate the indicator at OTU, Phylum, class, order, family, genus, and species level.

### 3.4 Alpha diversity analysis

The study examined alpha diversity through the analysis of the Chao1 and Shannon indexes. The calculations for Chao1 and Shannon index were conducted using QIIME version 1.9.1, while the OTUs rarefaction curve was generated using the R project ggplot2 package (version 2.2.1). Statistical analysis was carried out using Welch’s t method. The average Chao1 index for the Mock, APG, and N-APG samples were 737.43, 957.46, and 884.82, respectively (Figure 5A&B). Notably, no significant difference in the Chao1 index was observed within these groups (P>0.05, Figure 5C-E). On the other hand, the Shannon index of the Mock group was found to be higher compared to the APG and N-APG groups (Figure 5F-J). Specifically, the Shannon index for the Mock group ranged from 6.66 to 7.23, and its average was significantly higher than that of the APG group (Figure 5H).

**Figure 5.**
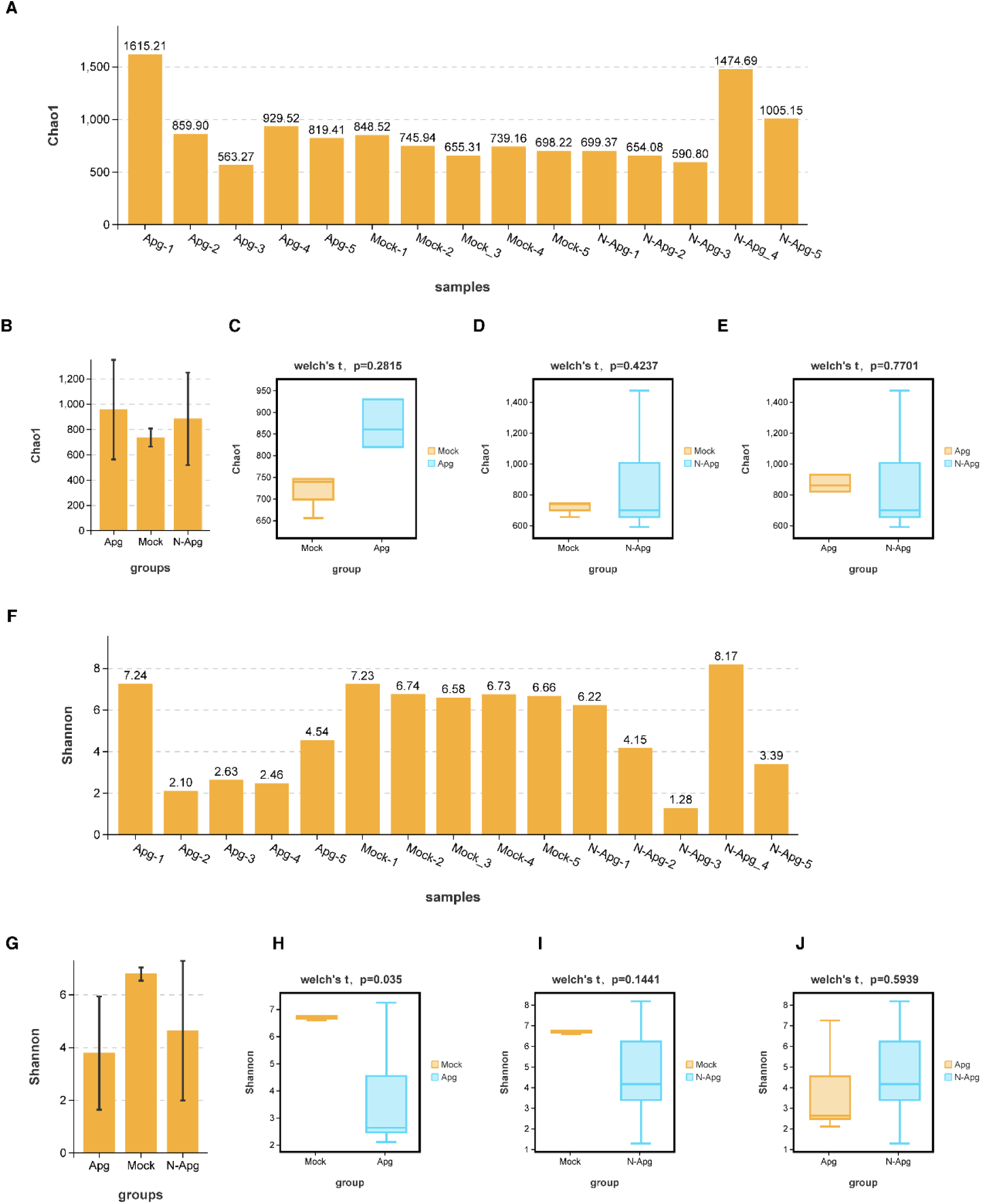
Alpha diversity analysis was conducted, resulting in the calculation of the Chao1 and Shannon indexes. (A) The Chao1 index was determined for each sample. (B) The average Chao1 index was calculated for three groups. (C-E) Statistical analysis was performed to compare the different groups. (F) The Shannon index was determined for each sample. (G) The average Shannon index was calculated for each group. (H-J) Statistical analysis was conducted to compare the different groups.

### 3.5 Beta diversity analysis

To analyze the bacterial communities among different groups, Bray-Curtis similarity was calculated, and Principal Coordinate Analysis (PCoA) was performed based on Bray-Curtis distances. This helped visualize the similarity of bacterial community structures among these groups (Figure 6A). The results showed that PCo1 and PCo2 accounted for 30.73% and 24.85% of the total variation, respectively. The samples displayed a distinct clustering in each group, and there was a partial crossover between the APG and N-APG groups. These findings suggested significant differences in the bacterial composition among the three groups. To further test this, the grouping information test was conducted, and unweighted-unifrac was selected for distance analysis. The Adonis (Permanova) test was calculated in the R project Vegan package (version 2.5.3). The Adonis test demonstrated that the beta diversity of the APG and N-APG group was significantly higher than that of the Mock group (Figure 6B and 6C, P=0.03 for APG v Mock, P=0.026 for N-APG v Mock). However, there was no significant difference between the beta diversity of APG and N-APG (P=0.28). Sample distance analysis showed that four of the mock samples were in one branch (Figure 6E), while among the APG or N-APG group, the samples showed different sample distances from each other (Figure 6E).

**Figure 6.**
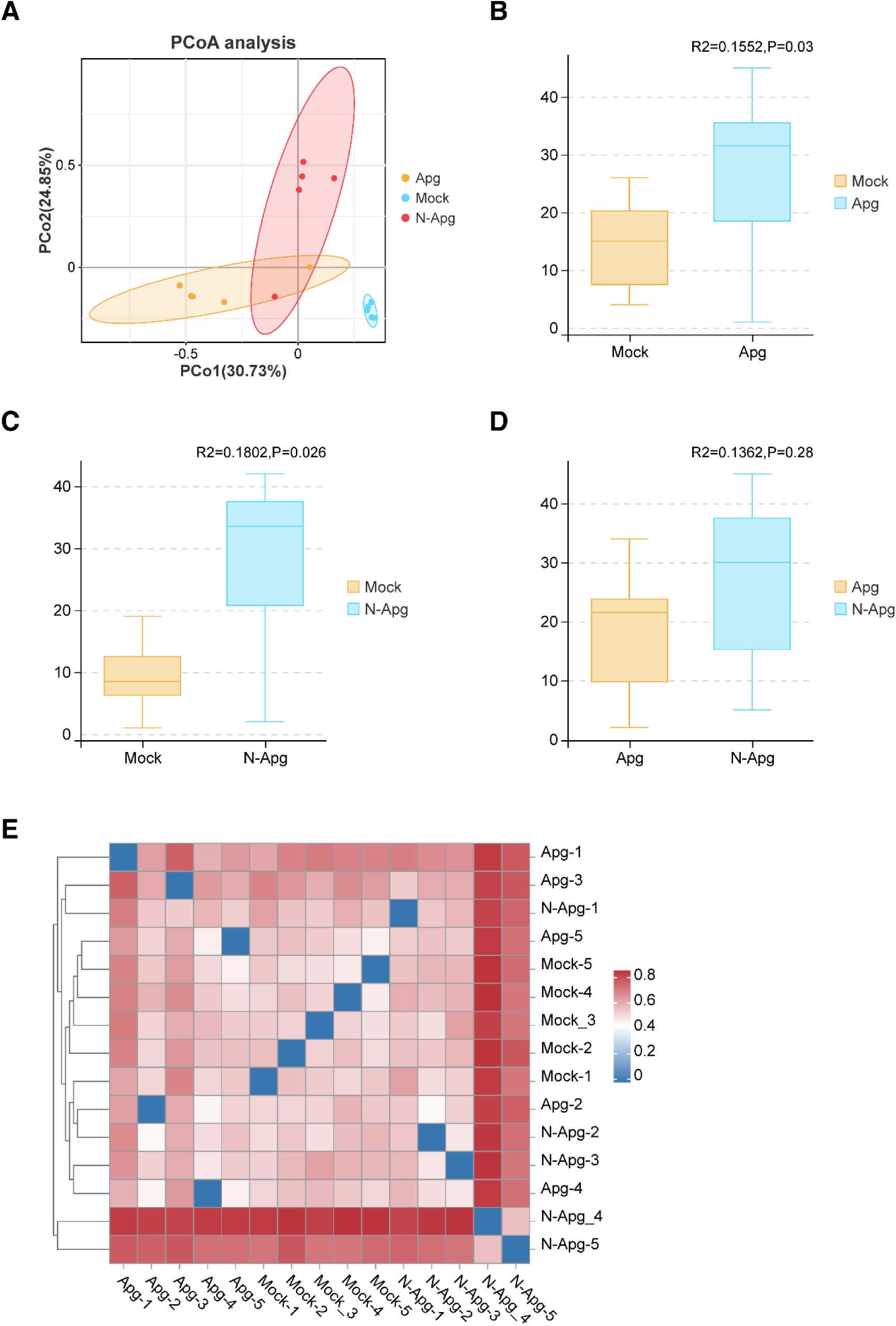
This study conducted beta diversity analysis using PCoA analysis for three groups and analyzed beta diversity between each group. We also created a species distance heatmap.

### 3.6 Function analysis

We utilized Tax4Fun (version 1.0) to conduct KEGG pathway analysis of the OTUs (Figure 7A). Our findings indicated that carbohydrate metabolism, membrane transport, translation, folding, sorting and degradation, nucleotide metabolism, replication, and repair, glycan biosynthesis, and metabolism pathway were downregulated in all samples of the APG group and 3/5 samples in the N-APG group, in comparison to the Mock group. Conversely, an upregulated pathway of amino acid metabolism, xenobiotics biodegradation and metabolism, signal transduction, cell motility, cell growth and death, lipid metabolism, metabolism of other amino acids, endocrine system, metabolism of terpenoids and polyketides, and biosynthesis of other secondary metabolites was detected in samples of the APG group and 3/5 samples in the N-APG group (Figure 7A). We also performed microbiome phenotype analysis using BugBase. Our findings indicated that the APG group demonstrated an upregulated level of contains mobile elements, stress-tolerant, potentially pathogenic, facultatively anaerobic, forms biofilms, and gram-negative (Figure 7B), while the APG and N-APG groups showed a decreased anerobic and gram-positive level (Figure 7B). Additionally, we utilized the FAPROTAX database (Functional Annotation of Prokaryotic Taxa) and associated software (version 1.0) to generate the ecological functional profiles of bacteria. Our analysis demonstrated that chicken samples with SHS had a higher abundance of animal parasites or symbionts (Figure 7C). In contrast, the bacterial community in the Mock group was associated with chemoheterotrophy, fermentation, animal parasites or symbionts, human gut, and mammal gut, etc (Figure 7C). Compared to the Mock group, most samples from the APG and N-APG groups showed an imbalanced bacteria community (Figure 7C).

**Figure 7.**
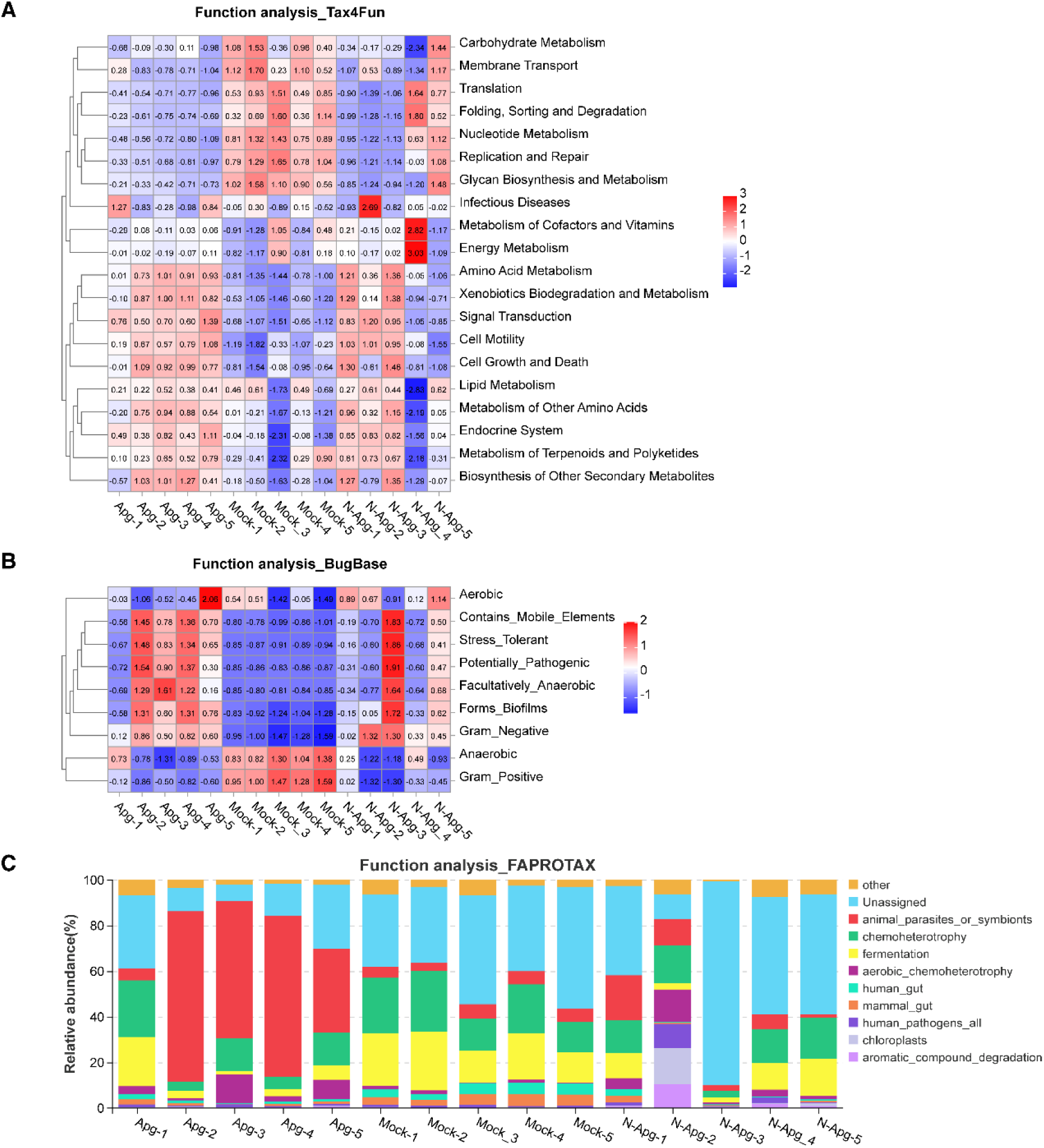
We used Tax4Fun (version 1.0) to infer KEGG pathway analysis of OTUs and conducted heat map analysis. Additionally, we classified microbiome phenotypes of bacteria using BugBase and generated ecological functional profiles of bacteria using FAPROTAX database (Functional Annotation of Prokaryotic Taxa) and associated software (version 1.0).

## 4 Discussion

Opportunistic pathogens are commonly present in healthy individuals but in low quantities. When the balance of the bacterial community is disrupted, these pathogens can proliferate and lead to local or systemic changes. In this study, 16S rRNA gene sequencing revealed the presence of APG in both APG PCR-positive and APG PCR-negative chickens, as well as SPF chickens.

Opportunistic pathogens can cause severe infections in individuals with compromised immune systems or microbiota. The symbiotic relationship between various microorganisms can hinder the colonization of opportunistic pathogens, preventing their expansion. However, these pathogens can initiate infection and expansion by inhibiting commensal bacteria, leading to changes in the microbiota’s structure and proportion. This disruption allows pathogenic bacteria to colonize and gain a growth advantage quickly. Recent studies suggest that *Staphylococcus chromogenes* plays a promoting role in infectious rhinitis caused by APG infection. It provides the necessary nutritional factor, nicotinamide adenine dinucleotide (NAD), for the growth of APG and accelerates its biosynthesis and release from host cells, promoting its survival and growth. Animal models have shown that antibiotics directed against *Staphylococcus chromogenes* can prevent APG infection (Wu et al., 2021).

PCR is a rapid technique for identifying pathogens based on their DNA. While conventional PCR can detect a range of DNA copies, real-time PCR is more sensitive and can detect lower levels of DNA. The intensity of the PCR signal is influenced by factors such as the copy number of the target gene, PCR reaction parameters, and cycling conditions. However, microbiota analysis has shown that PCR may not be sufficiently sensitive for pathogen identification. For instance, APG is an opportunistic pathogen of chickens that is commonly found in their infraorbital sinuses. Furthermore, microbiota sequencing has identified a significant number of animal pathogenic bacteria. It is worth noting that chickens harbor not only non-pathogenic bacteria, but also pathogenic bacteria that are capable of infecting humans.

Vaccines are highly effective tools in preventing the spread of infectious diseases (Zhang *et al*., 2015; Zhang *et al*., 2018a; Zhang *et al*., 2018b). Inactivated whole-cell and recombinant subunit protein vaccines are commonly used to protect against specific pathogens (Zhang et al., 2016; Zhang *et al*., 2015). It’s reported that double vaccination with an inactivated IC vaccine is more effective in preventing infectious coryza caused by APG serovar A, B, and C, compared to a single vaccination (Guo et al., 2022). Characterizing the pathogenicity of APG isolates is essential for developing an effective inactivated IC vaccine (Caballero-Garcia et al., 2022).

It is essential to acknowledge the limitations of this study. While conventional PCR can identify pathogens, it cannot quantify them. Future studies may consider using real-time PCR in conjunction with 16S rRNA sequencing to address this. It should also be noted that there are limitations to 16S rRNA sequencing, including the high cost of equipment compared to PCR testing, and its restriction to bacterial pathogens. Additionally, 16S rRNA sequencing may not effectively discriminate between certain species due to high sequence similarities. However, this can be resolved by altering the specific region of the 16S rRNA gene sequence (Fida *et al*., 2021). It is worth noting that 16S rRNA gene sequencing is based on PCR amplification, which can lead to the amplification of genes with high abundance and the omission of those with low abundance. Additionally, base pair mismatches may be introduced during the sequencing process.

To summarize, utilizing the sequencing of the 16S rRNA gene is a potent means of identifying bacterial pathogens. This technique applies to analyzing bacterial communities in samples and identifying new bacteria. In clinical laboratories, PCR is more commonly used than 16S rRNA sequencing. While it is the most widespread method, its sensitivity and accuracy do not match that of 16S rRNA sequencing. The signal intensity from conventional PCR is contingent upon the bacterial abundance. As sequencing identification technology becomes more accessible and cost-effective, it is likely more clinical microbiologists will adopt this method in their lab’s workflow.

## AUTHOR CONTRIBUTIONS

Conceptualization, planning, and management: GF Zhang. Laboratory experimental design: GF Zhang. Laboratory execution of experiments, and data acquisition, analysis, and interpretation: MN Li, N Wang, T Wang, S Zhang, Q Han, CC Zhang, YQ Shi, PZ Qiao, CL Man, T Feng, YY Li, XM You, Z Zhu, KJ Quan, TL Xu, GF Zhang. Drafting and revision of the manuscript: GF Zhang. Manuscript editing and approval: GF Zhang, TL Xu, KJ Quan, Z Zhu.

## ACKNOWLEDGMENTS

This study was supported by the Taishan Industrial Expert Programme (tscx202306107).

## DATA AVAILABILITY STATEMENT

Data that support the findings of this study are available from the corresponding author upon reasonable request.

---

**Supplemental Figure 1.**
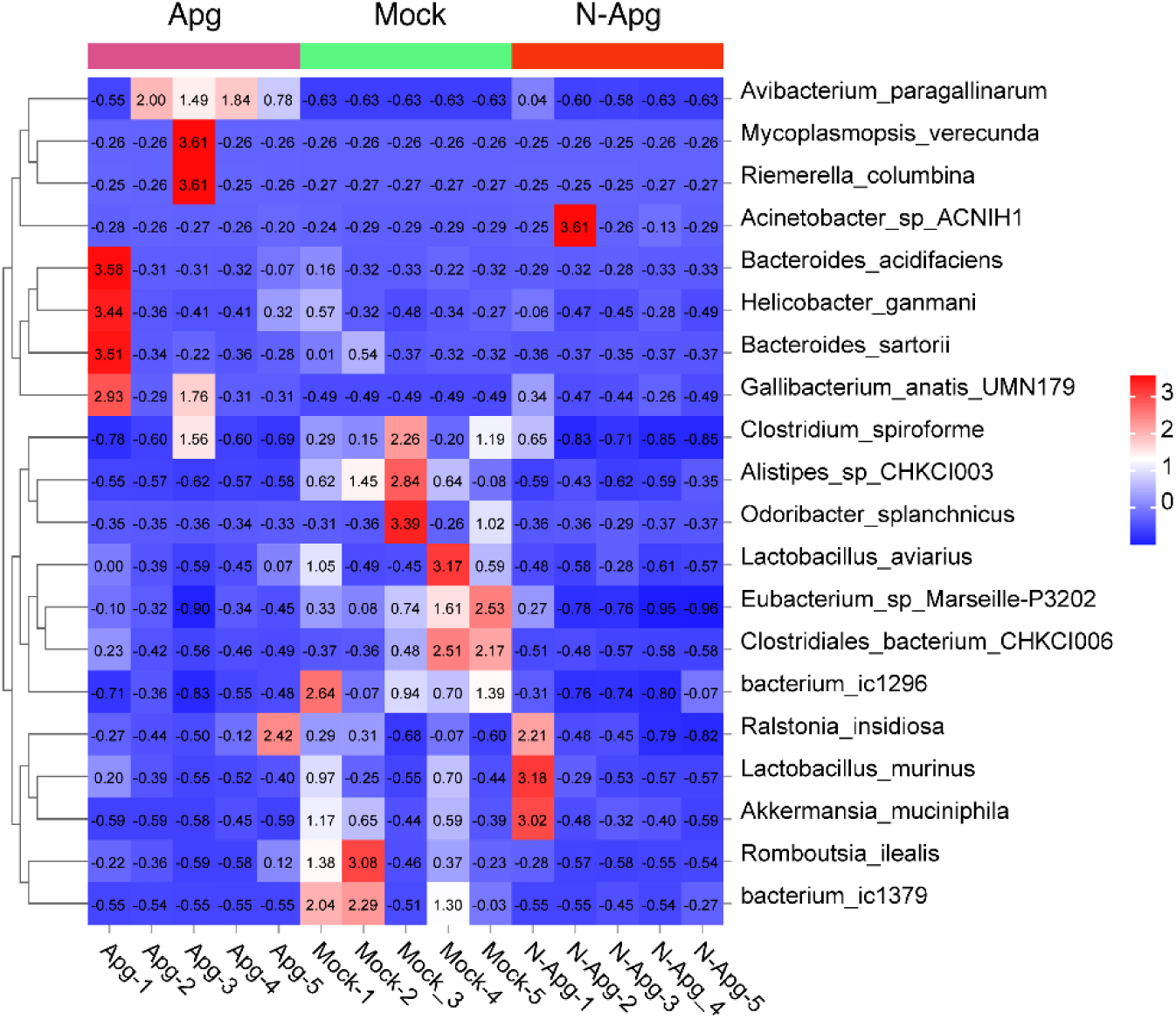
The heatmap of species identified in different samples.

**Supplemental Figure 2.**
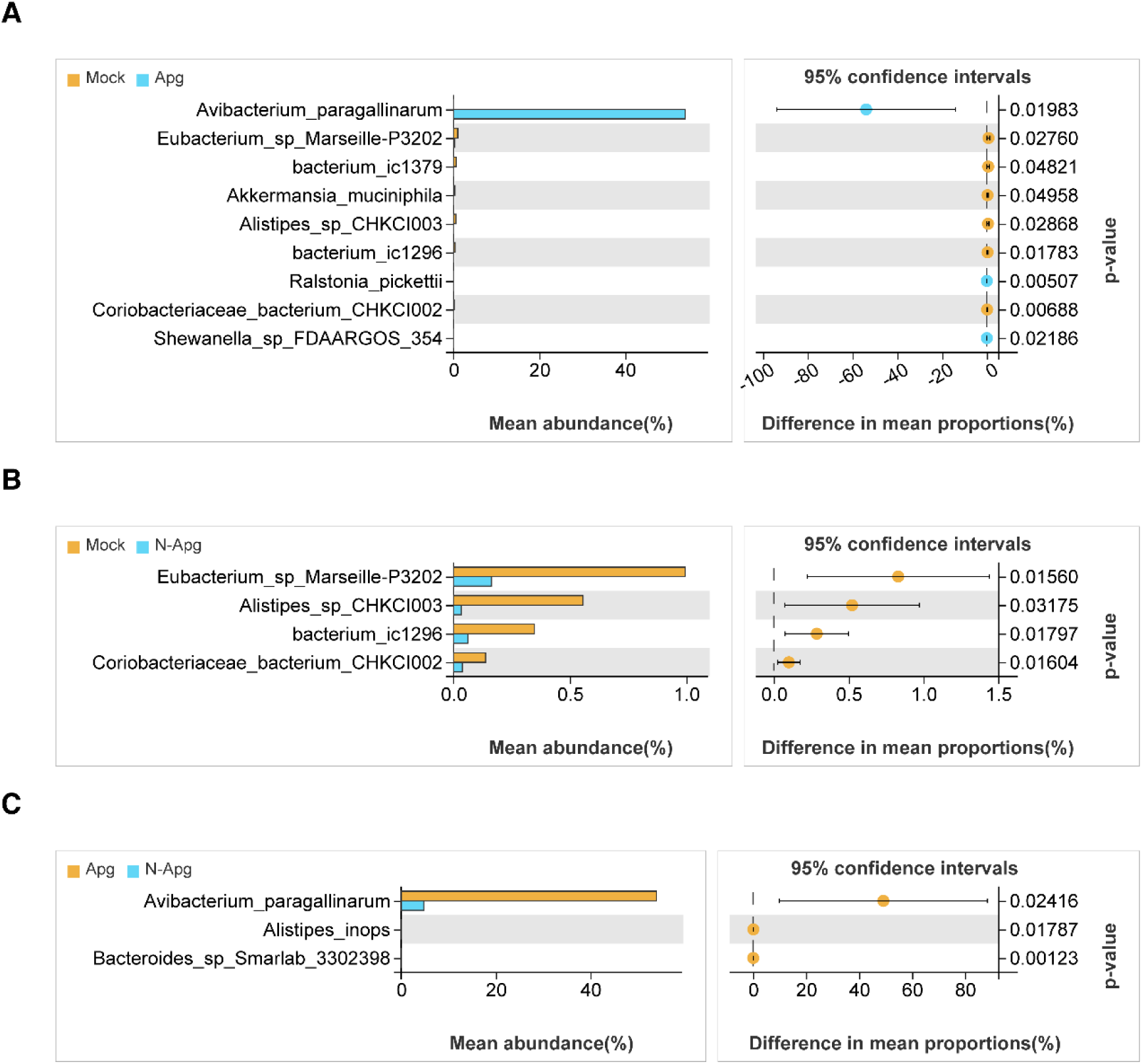
Indicator species analysis.

