## Supplemental Table 1 for "A detailed analysis of 16S rRNA gene sequencing and conventional PCR-based testing for the diagnosis of bacterial pathogens and discovery of novel bacteria"

| Species | Apg-1 | Apg-2 | Apg-3 |
|---|---|---|---|
| Avibacterium_paragallinarum | 2.6929 | 81.2631 | 65.4962 |
| Ralstonia_insidiosa | 1.4952 | 1.0337 | 0.8686 |
| Lactobacillus_aviarius | 1.5713 | 0.5637 | 0.0759 |
| Bacteroides_acidifaciens | 15.4287 | 0.1131 | 0.1095 |
| Acinetobacter_sp_ACNIH1 | 0.0172 | 0.0525 | 0.0336 |
| Eubacterium_sp_Marseille-P3202 | 0.428 | 0.3168 | 0.0336 |
| Romboutsia_ilealis | 0.2648 | 0.1625 | 0.0022 |
| Clostridium_spiroforme | 0.029 | 0.108 | 1.0324 |
| Lactobacillus_murinus | 0.3971 | 0.0946 | 0.0108 |
| Clostridiales_bacterium_CHKCI006 | 0.3491 | 0.0699 | 0.0076 |
| bacterium_ic1379 | 0 | 0.0051 | 0 |
| Akkermansia_muciniphila | 0.0036 | 0.0051 | 0.0054 |
| Alistipes_sp_CHKCI003 | 0.0236 | 0.0165 | 0 |
| Gallibacterium_anatis_UMN179 | 1.2894 | 0.0771 | 0.8513 |
| Mycoplasmopsis_verecunda | 0.0027 | 0.001 | 2.5343 |
| Riemerella_columbina | 0.0127 | 0.0051 | 2.4616 |
| Helicobacter_ganmani | 1.302 | 0.0463 | 0.0282 |
| bacterium_ic1296 | 0.0317 | 0.0915 | 0.0108 |
| Bacteroides_sartorii | 1.4607 | 0.0113 | 0.0575 |
| Odoribacter_splanchnicus | 0.0054 | 0.0051 | 0.0011 |
| Bacteroides_fragilis | 1.6239 | 0 | 0.0098 |
| Rothia_nasimurium | 0.0617 | 0.0031 | 0.0282 |
| Plesiomonas_shigelloides | 0.0054 | 0.0051 | 0 |
| Alistipes_finegoldii | 0.729 | 0.0555 | 0.0239 |
| Mycoplasmopsis_gallinacea | 9.00E-04 | 0.0031 | 1.3978 |
| bacterium_ic1391 | 0.0109 | 0.035 | 0 |
| Anoxybacillus_flavithermus | 9.00E-04 | 0 | 0 |
| Alistipes_inops | 0.1369 | 0.145 | 0.0098 |
| Ralstonia_pickettii | 0.0054 | 0.0082 | 0.0065 |
| Massiliomicrobiota_timonensis | 0 | 0.0051 | 0.0119 |
| Bacteroides_stercorirosoris | 0.904 | 0 | 0.0174 |
| Coriobacteriaceae_bacterium_CHKCI002 | 0.0571 | 0.0093 | 0.0011 |
| Paracoccus_yeei | 9.00E-04 | 0.0165 | 0.0076 |
| Ornithobacterium_rhinotracheale | 0.0073 | 0.0134 | 0.6539 |
| Enterococcus_cecorum | 0.1288 | 0.0134 | 0.0033 |
| Shewanella_sp_FDAARGOS_354 | 0.0027 | 0.0041 | 0.013 |
| Aerococcus_urinaeequi | 0.0444 | 0.0041 | 0.0076 |
| Bacteroidales_bacterium_RM8 | 9.00E-04 | 0 | 0 |
| Bacteroides_sp_Smarlab_3302398 | 0.0517 | 0.0401 | 0.0304 |
| Carnobacterium_inhibens_subsp_inhibens_DSM_13024 | 0.7299 | 0.001 | 0.0065 |
| Desulfovibrio_fairfieldensis | 0.437 | 0.0113 | 0.013 |
| Parabacteroides_goldsteinii | 0.6392 | 0.001 | 0 |
| Faecalibaculum_rodentium | 0.0018 | 0 | 0.0011 |
| Lactobacillus_coleohominis | 0 | 0 | 0.0033 |
| Comamonas_aquatica | 0 | 0 | 0 |
| Serratia_marcescens_WW4 | 0 | 0.0123 | 0.0043 |
| Lachnospiraceae_bacterium_28-4 | 0.0372 | 0.0113 | 0.0011 |
| Anaerotruncus_colihominis_DSM_17241 | 0.0036 | 0.001 | 0 |
| Prevotella_copri_DSM_18205 | 0.0997 | 0.0185 | 0.0065 |
| Methylobacterium_sp_PC3044 | 9.00E-04 | 0 | 0.0022 |
| Clostridiales_bacterium_CIEAF_020 | 0.1215 | 0.0041 | 0 |
| Streptococcus_respiraculi | 0.0018 | 0.0103 | 0.0011 |
| Sphingobacterium_faecium | 0.2711 | 0 | 0.0033 |
| Bifidobacterium_thermophilum_RBL67 | 0.0145 | 0.0031 | 0 |
| Bacteroides_vulgatus | 0.0036 | 0.0041 | 0.0065 |
| Macrococcus_caseolyticus | 0.0181 | 0.0226 | 0.0195 |
| Sphingomonas_paucimobilis | 0.0027 | 0 | 0.0011 |

| | | | |
|---|---|---|---|
| Muribaculum_intestinale | 0.0027 | 0.0041 | 0 |
| Corynebacterium_tuberculostearicum | 0.1324 | 0 | 0.0054 |
| Brachybacterium_paraconglomeratum | 9.00E-04 | 0.1265 | 0.0423 |
| Mycoplasmopsis_bovigenitalium | 0.2965 | 0.001 | 0 |
| Herbaspirillum_huttiense | 0.1877 | 0.0031 | 0.0011 |
| Staphylococcus_lentus | 9.00E-04 | 0.001 | 0 |
| Bacillus_thermoamylovorans | 0.0073 | 0 | 0.0011 |
| Deinococcus_antarcticus | 0.0508 | 0.0154 | 0 |
| bacterium_YE57 | 0.0018 | 0.0021 | 0 |
| Mycoplasmopsis_synoviae | 9.00E-04 | 0.001 | 0 |
| Streptococcus_pluranimalium | 0.1868 | 0 | 0 |
| Acinetobacter_gandensis | 0 | 0.001 | 0.0054 |
| Rothia_kristinae | 0.0018 | 0.001 | 0.0022 |
| Anaerofustis_sp_Marseille-P2832 | 0.0018 | 0.0031 | 0 |
| Anaerotignum_propionicum_DSM_1682 | 0 | 0 | 0 |
| Ihubacter_massiliensis | 0.0073 | 0.0051 | 0 |
| Acutalibacter_muris | 0.1786 | 0.0031 | 0.0065 |
| Nitrospira_japonica | 0 | 0 | 0 |
| Mycoplasmopsis_gallinarum | 0 | 0 | 0.2169 |
| Veillonella_magna | 0 | 0 | 0 |
| Gallibacterium_anatis | 0.1804 | 0.0103 | 0.0043 |
| Rhodobacteraceae_bacterium | 0.0018 | 0.0021 | 0 |
| Neisseria_sp_PPQC13-MTSKWH3-1 | 0 | 0 | 0.1941 |
| Corynebacterium_stationis | 0.0644 | 0 | 0.0065 |
| Succiniclasticum_ruminis | 9.00E-04 | 0 | 0 |
| Bacteroides_sp_Marseille-P3108 | 0 | 0.001 | 0 |
| Exiguobacterium_sp_AT1b | 0.0045 | 0 | 0 |
| Burkholderiales_bacterium_YL45 | 0 | 0 | 0.0011 |
| TM7_phylum_sp_oral_clone_CW040 | 0 | 0 | 0 |
| Sphingobacterium_multivorum | 0 | 0.001 | 0 |
| Aeriscardovia_aeriphila | 0.0154 | 0.0298 | 0.0022 |
| rumen_bacterium_enrichment_culture_clone_Y117 | 0 | 0 | 0 |
| Phascolarctobacterium_faecium | 0.0154 | 0.0195 | 0 |
| Diaphorobacter_polyhydroxybutyrativorans | 0.0335 | 0.0041 | 0.0087 |
| Lactobacillus_agilis | 0 | 0.0329 | 0 |
| Clostridium_merdae | 0 | 0 | 0 |
| Lactobacillus_ingluviei | 0 | 0.0494 | 0 |
| Fibrobacter_intestinalis | 0.0018 | 0 | 0.0011 |
| Pseudomonas_boreopolis | 0 | 0 | 0 |
| Fibrobacter_sp_UWB8 | 0 | 0 | 0 |
| Dysgonomonas_sp_HGC4 | 0 | 0 | 0.0065 |
| Helicobacter_hepaticus | 0 | 0 | 0.0076 |
| Lactobacillus_sp_30A | 0 | 0 | 0 |
| Other | 1.0625 | 0.3961 | 0.1781 |
| Unclassified | 64.5989 | 14.782 | 23.3972 |

| Apg-4 | Apg-5 | Mock-1 | Mock-2 | Mock_3 | Mock-4 | Mock-5 | N-Apg-1 | N-Apg-2 |
|---|---|---|---|---|---|---|---|---|
| 76.584 | 43.7621 | 0.015 | 0.0073 | 0.034 | 0.0401 | 0.0062 | 20.9645 | 1.1766 |
| 1.8867 | 8.7016 | 2.9995 | 3.0468 | 0.3965 | 2.0325 | 0.6151 | 8.1327 | 0.9224 |
| 0.4349 | 1.7275 | 4.1987 | 0.3251 | 0.4147 | 9.5678 | 3.0566 | 0.3567 | 0.1065 |
| 0.0494 | 1.0568 | 1.9411 | 0.0619 | 0.0364 | 0.4544 | 0.0697 | 0.1733 | 0.0626 |
| 0.0532 | 0.1536 | 0.0809 | 0 | 0 | 0 | 0.0042 | 0.0667 | 6.608 |
| 0.3086 | 0.2551 | 0.636 | 0.5163 | 0.8391 | 1.2676 | 1.7162 | 0.6089 | 0.0944 |
| 0.0104 | 0.5003 | 1.3889 | 2.5788 | 0.0922 | 0.6757 | 0.256 | 0.2222 | 0.0187 |
| 0.1082 | 0.067 | 0.4863 | 0.4289 | 1.3301 | 0.2805 | 0.8732 | 0.6434 | 0.0093 |
| 0.0275 | 0.0896 | 0.7998 | 0.1648 | 0.0073 | 0.6575 | 0.0645 | 1.9423 | 0.1449 |
| 0.0513 | 0.0394 | 0.0899 | 0.0956 | 0.4608 | 1.3377 | 1.1875 | 0.0311 | 0.0439 |
| 0.001 | 0 | 0.9965 | 1.0891 | 0.0158 | 0.7085 | 0.1977 | 0.0011 | 0 |
| 0.0522 | 0.0039 | 0.5951 | 0.4198 | 0.0534 | 0.4007 | 0.0718 | 1.219 | 0.0402 |
| 0.0152 | 0.0128 | 0.4014 | 0.672 | 1.124 | 0.4098 | 0.1748 | 0.0089 | 0.0607 |
| 0.0693 | 0.068 | 0 | 0 | 0 | 9.00E-04 | 0 | 0.3156 | 0.0093 |
| 0.0028 | 0.003 | 0 | 0 | 0 | 0 | 0 | 0.0056 | 0.0037 |
| 0.0104 | 0.0049 | 0 | 0 | 0 | 0 | 0 | 0.0122 | 0.0112 |
| 0.0304 | 0.2708 | 0.3535 | 0.0601 | 0.0073 | 0.0519 | 0.077 | 0.1456 | 0.0103 |
| 0.0598 | 0.0719 | 0.6121 | 0.1421 | 0.3177 | 0.2759 | 0.3944 | 0.1 | 0.0234 |
| 0.0038 | 0.0364 | 0.1428 | 0.3433 | 0 | 0.0209 | 0.0187 | 0.0033 | 0 |
| 0.0085 | 0.0148 | 0.021 | 0.0018 | 1.4344 | 0.0401 | 0.5287 | 0.0011 | 0.0028 |
| 0.0019 | 0.002 | 0 | 0 | 0 | 0 | 0 | 0.0056 | 0.0299 |
| 0.0712 | 0.002 | 0.001 | 0 | 0.0582 | 0 | 0 | 0.0945 | 1.1205 |
| 0.001 | 0.0039 | 0.002 | 0.0109 | 0.3953 | 0 | 0.0042 | 0.0111 | 0.0037 |
| 0.0532 | 0.0355 | 0.3375 | 9.00E-04 | 0.0182 | 0.0401 | 0.0885 | 0.0544 | 0.0037 |
| 0.0019 | 0.001 | 0 | 0 | 0 | 0 | 0 | 0 | 0.0028 |
| 0.018 | 0.0295 | 0.0789 | 0.0947 | 0.3832 | 0.1211 | 0.5266 | 0.03 | 9.00E-04 |
| 0.001 | 0 | 1.2411 | 0.0291 | 0.0036 | 0.0055 | 0.0062 | 0 | 0 |
| 0.1035 | 0.1477 | 0.2077 | 0.0401 | 0.017 | 0.2104 | 0.1259 | 0.0367 | 0 |
| 0.0076 | 0.0079 | 0.006 | 0 | 0 | 0 | 0 | 0.1333 | 0.015 |
| 0.001 | 0.001 | 0.2207 | 0.5536 | 0.0036 | 0.2732 | 0.0489 | 0 | 9.00E-04 |
| 0.001 | 0 | 0.0729 | 9.00E-04 | 0.0024 | 0.0273 | 0 | 0.0733 | 0 |
| 0.019 | 0.0069 | 0.1378 | 0.0719 | 0.1067 | 0.1466 | 0.2258 | 0.1089 | 9.00E-04 |
| 0.0142 | 0.0384 | 0.031 | 0.0036 | 0.0097 | 0 | 0 | 0.0344 | 0.6056 |
| 0.1272 | 0.0069 | 0 | 0 | 0 | 0 | 0 | 0.0378 | 0.0065 |
| 0.0038 | 0.003 | 0.0879 | 0.0319 | 0.023 | 0.0974 | 0.0021 | 0.1011 | 0.2813 |
| 0.0066 | 0.0059 | 0 | 0 | 0 | 0 | 0 | 0.0067 | 0.8401 |
| 0.0342 | 0.5959 | 0.0919 | 0 | 0 | 0 | 0 | 0.05 | 0.0392 |
| 0.001 | 0 | 0 | 0 | 0 | 0 | 0 | 0 | 0 |
| 0.0285 | 0.0522 | 0.1378 | 0.275 | 0.0036 | 0.112 | 0.0676 | 0.0011 | 0.0047 |
| 0.0047 | 0.0049 | 0 | 0 | 0 | 0 | 0 | 0.0011 | 0 |
| 0.0019 | 0.0059 | 0.0729 | 0 | 0 | 0.0191 | 0.0083 | 0.0767 | 0.0112 |
| 0.0019 | 0 | 0 | 0 | 0 | 0 | 0 | 0 | 0 |
| 0.0066 | 0 | 0.1707 | 0.1084 | 0.023 | 0.0838 | 0.0083 | 0.2333 | 9.00E-04 |
| 0 | 0 | 0.002 | 0.0046 | 0.5335 | 0.0046 | 0.0073 | 0 | 0 |
| 0 | 0.0374 | 0.0529 | 0 | 0 | 0.0255 | 0 | 0.0022 | 0.3934 |
| 0.0028 | 0.0394 | 0.1388 | 0.0146 | 0.023 | 0.0027 | 0.0052 | 0.0256 | 0.0028 |
| 0.0123 | 0.0049 | 0.0569 | 0.1393 | 0.0024 | 0.0619 | 0.0156 | 0.0033 | 0.1271 |
| 0 | 0 | 0.015 | 0.0291 | 0.1176 | 0.0355 | 0.2165 | 0.02 | 0 |
| 0.0133 | 0.1103 | 0.016 | 0 | 0.1103 | 9.00E-04 | 0.0021 | 0.0011 | 0.0215 |
| 0.0104 | 0.0295 | 0 | 0 | 0 | 0 | 0 | 0.0389 | 0.0178 |
| 0.001 | 0.002 | 0 | 0.051 | 0 | 0 | 0 | 0.1545 | 0.0075 |
| 0.0019 | 0 | 0.0459 | 9.00E-04 | 0 | 0.2705 | 0 | 0.0089 | 0 |
| 0 | 0.002 | 0 | 0 | 0 | 0 | 0 | 0.0022 | 0.0533 |
| 0.0076 | 0 | 0.2896 | 0.0027 | 0.0036 | 9.00E-04 | 0.0031 | 0 | 0 |
| 0.0057 | 0.0158 | 0.1068 | 0 | 0.023 | 0.0164 | 0.0219 | 0.1256 | 0.0019 |
| 0.0361 | 0 | 0 | 0.1266 | 0.0073 | 0.02 | 0.0021 | 0.0744 | 0 |
| 0 | 0.0148 | 0.007 | 0.0219 | 0.0061 | 0.0392 | 0 | 0 | 0.0159 |

| | | | | | | | | |
|---|---|---|---|---|---|---|---|---|
| 0.001 | 0 | 0 | 0 | 0 | 0 | 0 | 0.2711 | 0 |
| 0.0019 | 0.0817 | 0 | 0 | 0 | 0 | 0.0021 | 0.0956 | 0.0019 |
| 0.1301 | 0.002 | 0 | 0 | 0.0073 | 0 | 0 | 0 | 0.0019 |
| 0.0047 | 0.0049 | 0 | 0 | 0 | 0 | 0 | 0 | 0 |
| 0 | 0.001 | 0 | 0 | 0 | 9.00E-04 | 0 | 0 | 0.0019 |
| 0.0826 | 0.0049 | 0.027 | 0.0082 | 0.0012 | 0.01 | 0 | 0.1 | 0.0533 |
| 0.019 | 0 | 0.2556 | 0 | 0.0073 | 0.0046 | 0 | 0 | 0 |
| 0.0209 | 0.1024 | 0 | 0 | 0 | 0 | 0 | 0.1 | 0.0037 |
| 0.001 | 0.0276 | 0.022 | 0.0428 | 0.04 | 0.0382 | 0.076 | 0.0344 | 0 |
| 0.2754 | 0 | 0 | 0 | 0 | 0 | 0 | 0 | 9.00E-04 |
| 0 | 0.0256 | 0 | 0 | 0.0194 | 0 | 0 | 0.0022 | 0 |
| 0.0019 | 0.0207 | 0.001 | 0 | 0 | 0.0073 | 0.0031 | 0.0011 | 0.2243 |
| 0 | 0.2137 | 0 | 0 | 0.0242 | 0 | 0 | 0.0011 | 0 |
| 0 | 0 | 0.0609 | 0.0137 | 0 | 0.0446 | 0.0312 | 0.0789 | 0.0084 |
| 0.001 | 0 | 0 | 0 | 0 | 0 | 0 | 0 | 0 |
| 0.0066 | 0.0079 | 0.001 | 0.0291 | 0.0449 | 0.041 | 0.0739 | 0.0011 | 0.0168 |
| 0.0028 | 0.0059 | 0 | 9.00E-04 | 0 | 0.0246 | 0.0021 | 0 | 0.0047 |
| 0 | 0 | 0 | 0 | 0 | 0 | 0 | 0 | 0 |
| 0 | 0.001 | 0 | 0 | 0.0061 | 0 | 0 | 0 | 0 |
| 0 | 0.0039 | 0 | 0 | 0 | 0 | 0 | 0.21 | 0 |
| 0.001 | 0.001 | 0 | 0 | 0 | 0 | 0 | 0.0022 | 0 |
| 0.0665 | 0.0158 | 0 | 0.1129 | 0 | 0 | 0 | 0 | 0 |
| 0 | 0 | 0 | 0 | 0 | 0 | 0 | 0 | 0 |
| 0.0522 | 0.0039 | 0.02 | 0 | 0.0182 | 0 | 0 | 0.0033 | 0 |
| 0 | 0 | 0 | 0 | 0 | 0 | 0 | 0 | 0 |
| 0 | 0 | 0 | 0 | 0.0097 | 0 | 0 | 0.1111 | 0 |
| 0.0066 | 0 | 0 | 0.0209 | 0.0061 | 0 | 0.0104 | 0 | 0 |
| 0 | 0.001 | 0.016 | 0.02 | 0 | 0.0182 | 0 | 0.1256 | 0 |
| 0 | 0 | 0 | 0 | 0 | 0 | 0 | 0.05 | 0 |
| 0.0038 | 0.1566 | 0 | 0 | 0.0048 | 0 | 0 | 0 | 0 |
| 0 | 0.002 | 0 | 0 | 0 | 0 | 0 | 0.1111 | 0.0019 |
| 0 | 0 | 0 | 0 | 0 | 0 | 0 | 0 | 0 |
| 0.018 | 0.0158 | 0.012 | 0 | 0.0073 | 0.0282 | 0.0302 | 0 | 0.015 |
| 0.0019 | 0.0512 | 0.024 | 0.0018 | 0 | 0.0191 | 0.0114 | 0 | 9.00E-04 |
| 0 | 0.0414 | 0 | 0 | 0.0133 | 0 | 0 | 0.07 | 0 |
| 0 | 0 | 0 | 0 | 0 | 0 | 0 | 0.03 | 0 |
| 0 | 0.0611 | 0 | 0 | 0.0024 | 0 | 0 | 0 | 0 |
| 0 | 0 | 0.026 | 0 | 0 | 0 | 0 | 0 | 0 |
| 0 | 0 | 0.1108 | 0 | 0 | 0.0127 | 0.0052 | 0 | 0 |
| 0 | 0 | 0.025 | 0 | 0 | 0 | 0 | 0 | 0 |
| 0 | 0 | 0 | 0 | 0 | 0 | 0 | 0 | 0 |
| 0 | 0 | 0.03 | 0 | 0 | 0.0783 | 0 | 0 | 0 |
| 0 | 0 | 0.0469 | 0 | 0 | 0.0519 | 0.0125 | 0 | 0 |
| 0.2373 | 0.3036 | 0.5724 | 0.327 | 0.2 | 0.3022 | 0.0957 | 0.5656 | 0.3746 |
| 18.6942 | 40.7976 | 79.3631 | 87.8574 | 91.1499 | 79.4812 | 88.9475 | 61.5345 | 86.3316 |

| N-Apg-3 | N-Apg_4 | N-Apg-5 | Domain | Phylum | Class | Order | Family | Genus |
|---|---|---|---|---|---|---|---|---|
| 1.7877 | 0.062 | 0 | Bacteria | Proteobac | Gammapr | Pasteurella | Pasteurella | Avibacteriu |
| 1.0031 | 0.0898 | 0.01 | Bacteria | Proteobac | Gammapr | Burkholde | Burkholde | Ralstonia |
| 0.8397 | 0.0294 | 0.1129 | Bacteria | Firmicutes | Bacilli | Lactobacil | Lactobacil | Lactobacill |
| 0.2045 | 0 | 0 | Bacteria | Bacteroido | Bacteroidi | Bacteroida | Bacteroida | Bacteroide |
| 0.053 | 0.2808 | 0.01 | Bacteria | Proteobac | Gammapr | Pseudomo | Moraxella | Acinetoba |
| 0.1028 | 0.0098 | 0.0027 | Bacteria | Firmicutes | Clostridia | Lachnospi | Lachnospi | Lachnoclo |
| 0.0119 | 0.0294 | 0.0391 | Bacteria | Firmicutes | Clostridia | Peptostrep | Peptostrep | Rombouts |
| 0.0617 | 0 | 0 | Bacteria | Firmicutes | Bacilli | Erysipelotr | Erysipelato | Erysipelato |
| 0.0216 | 0 | 0 | Bacteria | Firmicutes | Bacilli | Lactobacil | Lactobacil | Lactobacill |
| 0.0032 | 0 | 0 | Bacteria | Firmicutes | Bacilli | Erysipelotr | Erysipelato | Erysipelato |
| 0.039 | 0.0049 | 0.1065 | Bacteria | Firmicutes | Clostridia | Oscillospir | Ruminoco | Faecalibac |
| 0.0931 | 0.0686 | 0.0036 | Bacteria | Verrucomi | Verrucomi | Verrucomi | Akkerman | Akkermans |
| 0 | 0.0098 | 0.0883 | Bacteria | Bacteroido | Bacteroidi | Bacteroida | Rikenellac | Alistipes |
| 0.0195 | 0.0865 | 0.0018 | Bacteria | Proteobac | Gammapr | Pasteurella | Pasteurella | Gallibacter |
| 0.0043 | 0 | 0 | Bacteria | Firmicutes | Bacilli | Mycoplasr | Mycoplasr | Mycoplasr |
| 0.0108 | 0 | 0 | Bacteria | Bacteroido | Bacteroidi | Flavobacte | Weeksella | Riemerella |
| 0.0162 | 0.0718 | 0.0046 | Bacteria | Campiloba | Campylob | Campylob | Helicobac | Helicobact |
| 0.026 | 0.0163 | 0.1429 | Bacteria | Firmicutes | Clostridia | Lachnospi | Lachnospi | Ruminoco |
| 0.0087 | 0 | 0 | Bacteria | Bacteroido | Bacteroidi | Bacteroida | Bacteroida | Bacteroide |
| 0.0303 | 0 | 0 | Bacteria | Bacteroido | Bacteroidi | Bacteroida | Marinifilac | Odoribacte |
| 0.0076 | 0 | 0 | Bacteria | Bacteroido | Bacteroidi | Bacteroida | Bacteroida | Bacteroide |
| 0.0206 | 0.0131 | 0 | Bacteria | Actinobac | Actinobac | Micrococc | Micrococc | Rothia |
| 0.0108 | 0.9582 | 0.0555 | Bacteria | Proteobac | Gammapr | Enterobac | Enterobac | Plesiomon |
| 0.0195 | 0 | 0 | Bacteria | Bacteroido | Bacteroidi | Bacteroida | Rikenellac | Alistipes |
| 0.0022 | 0 | 0 | Bacteria | Firmicutes | Bacilli | Mycoplasr | Mycoplasr | Mycoplasr |
| 0 | 0 | 0 | Bacteria | Firmicutes | Bacilli | Erysipelotr | Erysipelato | Erysipelato |
| 0.013 | 0 | 0 | Bacteria | Firmicutes | Bacilli | Bacillales | Bacillaceae | Anoxybaci |
| 0.0238 | 0 | 0 | Bacteria | Bacteroido | Bacteroidi | Bacteroida | Rikenellac | Alistipes |
| 0.0022 | 0.1632 | 0.7855 | Bacteria | Proteobac | Gammapr | Burkholde | Burkholde | Ralstonia |
| 0 | 0 | 0 | Bacteria | Firmicutes | Bacilli | Erysipelotr | Erysipelato | Erysipelato |
| 0.0022 | 0 | 0 | Bacteria | Bacteroido | Bacteroidi | Bacteroida | Bacteroida | Bacteroide |
| 0.0411 | 0 | 0.0373 | Bacteria | Actinobac | Coriobacte | Coriobacte | Eggerthell | CHKCI002 |
| 0.0097 | 0 | 0.1948 | Bacteria | Proteobac | Alphaprot | Rhodobac | Rhodobac | Paracoccu |
| 0.0184 | 0.0882 | 0 | Bacteria | Bacteroido | Bacteroidi | Flavobacte | Weeksella | Ornithoba |
| 0.0011 | 0.1257 | 0.0046 | Bacteria | Firmicutes | Bacilli | Lactobacil | Enterococ | Enterococ |
| 0.0195 | 0 | 9.00E-04 | Bacteria | Proteobac | Gammapr | Alteromor | Shewanell | Shewanella |
| 0.0022 | 0 | 9.00E-04 | Bacteria | Firmicutes | Bacilli | Lactobacil | Aerococca | Aerococcu |
| 0 | 0.8113 | 0.0291 | Bacteria | Bacteroido | Bacteroidi | Bacteroida | Rikenellac | Rikenellace |
| 0.0011 | 0 | 0 | Bacteria | Bacteroido | Bacteroidi | Bacteroida | Rikenellac | Alistipes |
| 0.0032 | 0 | 0 | Bacteria | Firmicutes | Bacilli | Lactobacil | Carnobact | Carnobact |
| 0.0011 | 0 | 0 | Bacteria | Desulfoba | Desulfovib | Desulfovib | Desulfovib | Desulfovib |
| 0 | 0 | 0 | Bacteria | Bacteroido | Bacteroidi | Bacteroida | Tannerella | Parabacter |
| 0 | 0.0016 | 0 | Bacteria | Firmicutes | Bacilli | Erysipelotr | Erysipelotr | Faecalibac |
| 0 | 0.031 | 0.0046 | Bacteria | Firmicutes | Bacilli | Lactobacil | Lactobacil | Lactobacill |
| 0.0065 | 0 | 0 | Bacteria | Proteobac | Gammapr | Burkholde | Comamor | Comamon |
| 0.0108 | 0.0131 | 0.1939 | Bacteria | Proteobac | Gammapr | Enterobac | Yersiniace | Serratia |
| 0.0032 | 0 | 0 | Bacteria | Firmicutes | Clostridia | Lachnospi | Lachnospi | Blautia |
| 0 | 0 | 0 | Bacteria | Firmicutes | Clostridia | Oscillospir | Ruminoco | Anaerotru |
| 0 | 0 | 0 | Bacteria | Bacteroido | Bacteroidi | Bacteroida | Prevotella | Prevotella |
| 0 | 0.0996 | 0.1675 | Bacteria | Proteobac | Alphaprot | Rhizobiale | Beijerincki | Methyloba |
| 0 | 0 | 0 | Bacteria | Firmicutes | Clostridia | Lachnospi | Lachnospi | Lachnospi |
| 0 | 0 | 0 | Bacteria | Firmicutes | Bacilli | Lactobacil | Streptococ | Streptococ |
| 0 | 0 | 0 | Bacteria | Bacteroido | Bacteroidi | Sphingoba | Sphingoba | Sphingoba |
| 0.0065 | 0 | 0 | Bacteria | Actinobac | Actinobac | Bifidobact | Bifidobact | Bifidobacte |
| 0 | 0 | 0 | Bacteria | Bacteroido | Bacteroidi | Bacteroida | Bacteroida | Bacteroide |
| 0 | 0 | 0 | Bacteria | Firmicutes | Bacilli | Staphyloc | Staphyloc | Macrococ |
| 0 | 0.0212 | 0.1966 | Bacteria | Proteobac | Alphaprot | Sphingom | Sphingom | Sphingom |

| | | | | | | | | |
|---|---|---|---|---|---|---|---|---|
| 0.0454 | 0 | 0 | Bacteria | Bacteroid | Bacteroidi | Bacteroida | Muribacul | Muribacul |
| 0.0022 | 0 | 0 | Bacteria | Actinobac | Actinobac | Corynebac | Corynebac | Corynebac |
| 0.0011 | 0.0016 | 0 | Bacteria | Actinobac | Actinobac | Micrococc | Dermabac | Brachybac |
| 0 | 0 | 0 | Bacteria | Firmicutes | Bacilli | Mycoplasr | Mycoplasr | Mycoplasr |
| 0.0011 | 0.0016 | 0.1065 | Bacteria | Proteobac | Gammapr | Burkholde | Oxalobact | Herbaspiri |
| 0.0119 | 0 | 0 | Bacteria | Firmicutes | Bacilli | Staphyloc | Staphyloc | Staphyloc |
| 0.0043 | 0 | 0 | Bacteria | Firmicutes | Bacilli | Bacillales | Bacillacea | Bacillus |
| 0.0022 | 0 | 0 | Bacteria | Deinococc | Deinococc | Deinococc | Deinococc | Deinococc |
| 0 | 0 | 0 | Bacteria | Firmicutes | Clostridia | Christense | Christense | Christense |
| 0.0022 | 0 | 0 | Bacteria | Firmicutes | Bacilli | Mycoplasr | Mycoplasr | Mycoplasr |
| 0 | 0.0408 | 0 | Bacteria | Firmicutes | Bacilli | Lactobacil | Streptoco | Streptoco |
| 0.0022 | 0.0016 | 0 | Bacteria | Proteobac | Gammapr | Pseudomc | Moraxella | Acinetoba |
| 0.0032 | 0 | 0 | Bacteria | Actinobac | Actinobac | Micrococc | Micrococc | Kocuria |
| 0 | 0 | 0 | Bacteria | Firmicutes | Clostridia | Eubacteria | Anaerofus | Anaerofus |
| 0 | 0.2383 | 9.00E-04 | Bacteria | Firmicutes | Clostridia | Lachnospi | Lachnospi | Anaerostig |
| 0 | 0 | 0 | Bacteria | Firmicutes | Clostridia | Peptostrep | Anaerovor | Eubacteriu |
| 0.0011 | 0 | 0 | Bacteria | Firmicutes | Clostridia | Oscillospir | Ruminoco | Incertae_S |
| 0 | 0.018 | 0.2075 | Bacteria | Nitrospiro | Nitrospiria | Nitrospiral | Nitrospira | Nitrospira |
| 0 | 0 | 0 | Bacteria | Firmicutes | Bacilli | Mycoplasr | Mycoplasr | Mycoplasr |
| 0 | 0 | 0 | Bacteria | Firmicutes | Negativicu | Veillonella | Veillonella | Veillonella |
| 0.0011 | 0 | 0 | Bacteria | Proteobac | Gammapr | Pasteurella | Pasteurella | Gallibacter |
| 0 | 0 | 0 | Bacteria | Proteobac | Alphaprot | Rhodobac | Rhodobac | Roseovariu |
| 0.0022 | 0 | 0 | Bacteria | Proteobac | Gammapr | Burkholde | Neisseriac | Neisseria |
| 0 | 0.0261 | 0 | Bacteria | Actinobac | Actinobac | Corynebac | Corynebac | Corynebac |
| 0 | 0.1861 | 0.0073 | Bacteria | Firmicutes | Negativicu | Acidaminc | Acidaminc | Succiniclas |
| 0.0638 | 0 | 0 | Bacteria | Bacteroid | Bacteroidi | Bacteroida | Bacteroida | Bacteroide |
| 0 | 0.1273 | 0.0082 | Bacteria | Firmicutes | Bacilli | Exiguobac | Exiguobac | Exiguobac |
| 0 | 0 | 0 | Bacteria | Proteobac | Gammapr | Burkholde | Sutterellac | Parasutter |
| 0 | 0.1257 | 0.0046 | Bacteria | Patescibac | Saccharim | Saccharim | Saccharim | Candidatu |
| 0 | 0 | 9.00E-04 | Bacteria | Bacteroid | Bacteroidi | Sphingoba | Sphingoba | Sphingoba |
| 0 | 0 | 0 | Bacteria | Actinobac | Actinobac | Bifidobact | Bifidobact | Aeriscardc |
| 0 | 0.1567 | 0.0055 | Bacteria | Firmicutes | Clostridia | Lachnospi | Lachnospi | Oribacteriu |
| 0 | 0 | 0 | Bacteria | Firmicutes | Negativicu | Acidaminc | Acidaminc | Phascolarc |
| 0.0022 | 0 | 0 | Bacteria | Proteobac | Gammapr | Burkholde | Comamor | Diaphorob |
| 0 | 0 | 0 | Bacteria | Firmicutes | Bacilli | Lactobacil | Lactobacil | Lactobacill |
| 0 | 0.111 | 0 | Bacteria | Firmicutes | Clostridia | Oscillospir | Ruminoco | Incertae_S |
| 0.026 | 0 | 0 | Bacteria | Firmicutes | Bacilli | Lactobacil | Lactobacil | Lactobacill |
| 0.0011 | 0.0996 | 0.0027 | Bacteria | Fibrobacte | Fibrobacte | Fibrobacte | Fibrobacte | Fibrobacte |
| 0 | 0 | 0 | Bacteria | Proteobac | Gammapr | Xanthomc | Xanthomc | Xylella |
| 0 | 0.1012 | 0 | Bacteria | Fibrobacte | Fibrobacte | Fibrobacte | Fibrobacte | Fibrobacte |
| 0 | 0.1159 | 0.0018 | Bacteria | Bacteroid | Bacteroidi | Bacteroida | Dysgonon | Dysgonom |
| 0 | 0 | 0 | Bacteria | Campiloba | Campylob | Campylob | Helicobac | Helicobact |
| 0 | 0 | 0 | Bacteria | Firmicutes | Bacilli | Lactobacil | Lactobacil | Lactobacill |
| 0.1765 | 1.9068 | 0.1144 | | | | | | |
| 95.0871 | 93.6563 | 97.3458 | | | | | | |

ccus_gauvreauii_group

acterium-Methylorubrum
raceae_NK4A136_group

nellaceae_R-7_group
